# Genome-wide targets of selection: female response to experimental removal of sexual selection in *Drosophila melanogaster*

**DOI:** 10.1101/000240

**Authors:** Paolo Innocenti, Ilona Flis, Edward H. Morrow

## Abstract

Despite the common assumption that promiscuity should in general be favored in males, but not in females, to date there is no consensus on the general impact of multiple mating on female fitness. Notably, very little is known about the genetic and physiological features underlying the female response to sexual selection pressures. By combining an experimental evolution approach with genomic techniques, we investigated the effects of single and multiple matings on female fecundity and gene expression. We experimentally manipulated the mating system in replicate populations of *Drosophila melanogaster* by removing sexual selection, with the aim of testing differences in short term post-mating effects of females evolved under different mating strategies. We show that monogamous females suffer decreased fecundity, a decrease that was partially recovered by experimentally reversing the selection pressure back to the ancestral promiscuous state. The post-mating gene expression profiles of monogamous females differ significantly from promiscuous females, involving 9% of the genes tested. These transcripts are active in several tissues, mainly ovaries, neural tissues and midgut, and are involved in metabolic processes, reproduction and signaling pathways. Our results demonstrate how the female post-mating response can evolve under different mating systems, and provide novel insights into the genes targeted by sexual selection in females, by identifying a list of candidate genes responsible for the decrease in female fecundity in the absence of promiscuity.

## Introduction

The evolution of mating strategies, and in particular of female polyandry, has attracted a great deal of attention in recent decades (Andersson, 1994; Birkhead, 2000; Shuster & Wade, 2003). The debate on the adaptive significance of female multiple mating stems from the common assumption that males exhibit a stronger positive covariance between promiscuity and reproductive success than females. In other words, males may gain more offspring by repeated matings than females, even though both sexes have the same average numbers of matings, mates and offspring (Shuster & Wade, 2003). Polyandry is also assumed to carry costs in terms of time and energy for additional matings (Jormalainen *et al.*, 2001; Wedell *et al.*, 2006) or physical injury (Boeuf & Mesnick, 1991; Hurst *et al.*, 1995), as well as an increased risk of predation (Wing, 1988) and infection during copulation (Hurst *et al.*, 1995).

Empirical studies however, show that in the vast majority of species, females often mate with more than one male (Ridley, 1988; Andersson, 1994; Simmons, 2001). Theoretically, female polyandry can be promoted by selection if males provide resource benefits, through the ejaculate (Thornhill, 1976; Boggs & Gilbert, 1979; Turner & Anderson, 1983) or through additional paternal care (Stacey, 1982; Davies, 1985), or if some males do not provide viable sperm or insufficient ejaculate to fertilize the ova (Gibson & Jewell, 1982; Gromko *et al.*, 1984). It has also be proposed as a strategy to reduce sexual harassment (Svärd & Wiklund, 1986). Moreover, there could be indirect genetic benefits by acquiring ‘good genes’, compatible genes or in producing genetically diverse progeny or promoting sperm competition (Yasui, 1998). Finally, multiple mating can be non-adaptive for females in the presence of strong selection for multiple mating on males coupled with a strong intersexual genetic correlation for mating propensity (Halliday & Arnold, 1987).

Recently, experimental evolution studies have been an increasingly popular approach to evaluate the effect of different mating systems on male or female fitness (Edward *et al.*, 2010), predominantly within the framework of sexually antagonistic co-evolution. Under a promiscuous mating system, where the fitness values of an individual and its mate are not perfectly correlated, pre-copulatory and post-copulatory intrasexual competition are expected to result in the evolution of traits that increase the reproductive success of members of one sex at the expense of the other, in a co-evolutionary arms race called interlocus sexual conflict (Parker, 1979). An eminent example of harm induced by males to females in an attempt to maximize their mating rate and fertilization success is represented by *Drosophila melanogaster*, in which courtship and transfer of seminal fluid are known to increase female mortality rate and decrease lifetime reproductive success while increasing male competitive abilities (Chapman *et al.*, 1995).

Holland and Rice (1999) were the first to investigate the change in female reproductive success in populations of *D. melanogaster* using an experimental evolution design where sexual selection was removed by enforcing monogamy and random mating assignment. They found that monogamous populations had greater net reproductive rate than (promiscuous) controls, while fecundity of monogamous females was reduced after mating with ancestral (promiscuous) males (Holland & Rice, 1999). After this seminal paper, several other studies employed a similar methodology in different taxa (Hosken *et al.*, 2001, 2009; Martin & Hosken, 2003; Crudgington *et al.*, 2005; Tilszer *et al.*, 2006; Bacigalupe *et al.*, 2007; Fricke & Arnqvist, 2007; LaMunyon *et al.*, 2007; Simmons & Garcia Gonzalez, 2008; Gay *et al.*, 2009; Maklakov *et al.*, 2009), briefly reviewed by Edward *et al.* (2010), with some degree of variation in experimental design and outcome.

Regardless of the adaptive significance of female polyandry, the genetic basis of the fitness components that depend on different mating strategies is a key aspect, which has so far received little attention. In other words, very little information is available about the characteristics and identity of the genes that respond to an alteration of sexual selection (i.e. the targets of selection, but see Immonen & Ritchie, 2011). With modern genomic techniques, it is possible to scan whole genomes and transcriptomes and associate them with the corresponding phenotypes. Coupling experimental evolution with genome sequencing or transcriptome profiling is a very recent and successful approach (Burke *et al.*, 2010; Zhou *et al.*, 2011), in that it experimentally magnifies the variation in the trait of interest and produces a greater resolving power in identifying structure of molecular networks and adaptive processes (Turner *et al.*, 2011). However, these methods to our knowledge have not been applied to sexual selection studies so far.

Conversely, other aspects of the fruit fly reproductive biology are much better known. In recent years, considerable quantities of data have been collected on the female physiological changes associated with the shifts in female mating status. Molecular and genomics techniques have been employed to investigate the effects of mating in *D. melanogaster* (Lawniczak & Begun, 2004; McGraw *et al.*, 2004, 2008; Mack *et al.*, 2006; Innocenti & Morrow, 2009). In particular, several detailed studies have focused on seminal fluid components on female post-mating physiology, leading to the identification of several seminal fluid proteins (SFP) and to the isolation of their effect in females (reviewed in Ravi Ram & Wolfner, 2007; Avila *et al.*, 2011), including the characterization of sex-peptide and its receptor (Domanitskaya *et al.*, 2007; Yapici *et al.*, 2008).

Here, we integrate these approaches to investigate the evolutionary response of populations experiencing differing sexual selection pressures, at both a phenotypic and genomic level, allowing a direct comparison between the two. We begin by using experimental evolution to evaluate the effects of the removal of components of sexual selection in a laboratory-adapted population of *D. melanogaster.* The effects of enforced monogamy are then investigated both in terms of differences in female reproductive output and in female post-mating response, measured as genome-wide gene expression profiles. In addition, for populations that have evolved under enforced monogamy we subsequently reverse the selection pressure back to the ancestral promiscuous state and again investigate how reproductive output is affected. We take an exploratory approach to investigate the characteristics and biology of those transcripts identified as being influenced by the experimental selection regimes, with the ultimate aim of understanding in more detail which biological processes in females are associated with evolutionary changes in mating system.

## Materials and Methods

### Fly stocks

All flies used to constitute the experimental evolution lines were derived from a large outbred wild-type population of *D. melanogaster* (LH_M_) that had been maintained under the same rearing protocol for over 400 non-overlapping generations (for a detailed description, see Rice *et al.*, 2005). The population is maintained in a set of 56 vials at a large size (1792 adults) under competitive conditions and at moderate larval density in standard rearing environment: 25°C, cornmeal/molasses/yeast/agar medium, 12h:12 h light/dark cycle, 16 individuals of each sex per vial (25 mm × 95 mm) with a 14 day generation cycle. We applied the same culturing condition to our experimental lines, unless otherwise specified.

### Experimental evolution lines

In March 2008, a replicate of the ancestral LH_M_ population was obtained by allowing females to lay eggs for 18 h (Day 0). On the day of emergence, Day 10, we collected 384 virgin males and 384 virgin females from the base population, and randomly assigned them to 2 treatments, each constituted by 4 replicate populations of 96 individuals, and stored separately by sex. On Day 13, males and females were placed together in fresh vials (16 pairs per vials, three vials per population) with 6 mg fresh yeast and allowed to mate. In one treatment (hereafter referred to as “monogamous treatment”), males were removed after 1 h under brief CO_2_ anesthesia and discarded. During this window of time, in our LH_M_ population virtually all the sexually mature and healthy females mate, but none of the females mate twice, due to their refractory period. We performed a preliminary study using time-lapse photography (see Kuijper & Morrow, 2009) to confirm this pattern: we placed 25 vials containing 10 virgin males and 10 virgin females in an incubator under standard conditions and monitored their activity for 12 h. The results show a peak in mating activity (often 10 pairs simultaneously) between 10 and 30 minutes, followed by a long (>1 h) refractory period, after which females start re-mating (Fig. S8). In the other treatment (hereafter “promiscuous treatment”), males were left in the vials with the females and allowed to mate further. On day 14 (i.e. Day 0 of the following generation), the flies were transferred to fresh vials to oviposit for 18 h. The following day (Day 1), eggs were counted and those exceeding 150 were removed to ensure a uniform larval environment On Day 10, 96 individuals per replicate population were collected as virgin (48 males and 48 females) and the same culturing conditions described above were applied every generation.

### Body size

After 30 generations of experimental evolution we harvested 40 males and 40 females from each replicate population to assess whether there had been a change in body size. A single wing was removed from each individual, mounted on a slide using transparent tape and photographed using bright field-illumination (x40 magnification). Length was measured using the straight line tool in ImageJ (Rasband, 1997), from the intersection of the anterior cross vein and longitudinal vein 3 (L3) to the intersection of L3 with the distal wing margin (Partridge *et al.*, 1987).

### Female fecundity

The effect of the treatment on female fecundity was assayed with a factorial design in four different trials, after 30, 31, 50 and 58 generations of experimental evolution, with slightly different experimental designs (trials 1-4 respectively, see below). For each trial, the following protocol was applied: on day 14 of the chosen generation, a replicate of the experimental lines were obtained, by allowing flies to oviposit for an additional 24 h in fresh vials. The populations obtained were cultured with standard protocol (i.e. as in the promiscuous treatment) for a generation to remove parental effects. On Day 10 of the following generation, 160 females and 160 males for each treatment and replicate were collected as virgins and stored separately (10 vials of males and 10 vials of females for each of the 8 experimental lines). On Day 13, half of the females from each experimental line (5 vials) were crossed to males from the same experimental line, and the other half (5 vials) were crossed to males from a single replicate population of the other treatment (females from replicate 1 of the monogamous treatment were mated to males from replicate 1 of the promiscuous treatment, replicate 2 of the monogamous treatment was paired with replicate 2 of the promiscuous treatment, and so on), and allowed to mate in fresh vials containing 6 mg live yeast. At this stage, the trials differed in their design, as follows.

In trial 1, males were removed and discarded the following day (after 30 h, Day 14), while individual females were transferred in oviposition test tubes, and allowed to oviposit for 18 hours, corresponding to the window of time in which eggs laid by females in the experimental populations were retained for the next generation. Females were then discarded, the tubes refrigerated for 24 hours and the eggs counted. In trial 2, the protocol employed was identical to the one described for trial 1, except the males were removed and discarded after 1 h, allowing females to mate only once.

In trial 3, after 1 h, all the males were removed and discarded. Females were allowed to oviposit, and were transferred every 12 h (at 9:00 and at 21:00) to a fresh vial for four days (6 times, 7 time-points) to avoid excessive larval density, then discarded. When the new generation emerged, progeny were counted. In trial 4, the protocol employed was very similar to the one described for trial 3, with the following differences: after crossing target males and females, males were not removed from the vials; also, during the four days of oviposition the flies were transferred every 6 h during the daylight hours (9:00, 15:00 and 21:00; 9 times, 10 time-points).

In summary, we obtained measures of fecundity of females in our experimental lines under four different conditions: after a single mating and after being continuously exposed to males, during the whole period in which they were allowed to oviposit in our selection regime, or during a longer timeframe, to account for potential shifts in the resource that females allocate to eggs over time.

### Reversed selection lines

After 95 generations of experimental evolution a third treatment was established using surplus flies harvested from each of the 4 monogamous populations. In this new treatment, the rearing protocol was identical to that for the promiscuous treatment, and therefore flies in these populations experienced a reversal of the selection pressure from a monogamous to a promiscuous mating system (hereafter refereed to as the MP treatment). After a further 25 generations of experimental evolution (generation 120 in total) the populations were cultured again with standard protocol for a single generation to remove parental effects, then an assay of female fitness from all replicate populations and treatments was performed (n = 53–64 individual females per population), using the same protocol employed for trial 1 (see above).

### Microarray data

After 46 generations, on day 14, replicates of the experimental lines were obtained, by allowing flies to oviposit for additional 24 h in fresh vials. The populations obtained were cultured with standard protocol (i.e. as the promiscuous treatment) for a generation to remove parental effects. On Day 10 of the second generation, 64 females and 64 males for each treatment and replicate were collected as virgins and stored separately (4 vials of males and 4 vials of females for each of the 8 experimental lines). On Day 13, half of the females from each experimental line (2 vials) were crossed to males from the same experimental line, and the other half (2 vials) were crossed to males from a single replicate population of the other treatment, and allowed to mate in fresh vials containing 6 mg live yeast After 1 h, all the males were removed and discarded, while the females were randomly divided in two groups of 8 flies under brief CO_2_ anesthesia, to be used as a main sample and its backup. After 6 hours, the females were flash-frozen in liquid nitrogen and stored at −80°C for no more than four days until RNA extraction. Hence, for each of the replicate population we collected 4 independent samples of eight females, two samples of females mated to males of the same replicate population, and two samples of females mated to males of a single replicate population in the other treatment, giving a total of 32 samples. Total RNA was extracted independently from each sample using Trizol (Invitrogen, Carlsbad, CA, USA) and purified with an RNeasy Mini Kit (Qiagen, Hilden, Germany). RNA quality and quantity was assessed with an Agilent Bioanalyzer (Agilent Technologies, Santa Clara, CA, USA). The RNA samples were prepared and hybridized to Affymetrix Drosophila GeneChip 2.0 microarrays at the Uppsala Array Platform (Uppsala, Sweden) following manufacturer’s instructions. The arrays were scanned in two batches of 16, balanced for replicate population of origin and replicate population of origin of the males to which they were mated.

### Statistical analysis

All statistical analyses were run in the R environment (version 2.11.1 for most analyses, version 3.0.1 for body size and reversed experimental evolution assay, available at www.r-projectorg R Development Core Team, 2009).

Male and female body size was analyzed using a full factorial linear model (lm function; mating system and sex as fixed effects) using within replicate means to avoid pseudoreplication (n = 36).

Female fecundity data for each trial were analyzed using linear models (lm function). In all cases, amount of eggs or progeny produced was averaged within replicate population and summed across time points (for trials 3 and 4), to avoid pseudoreplication. We fitted the following model to each dataset:

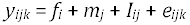

with *i* = {1,2}; j = {1,2}; k = {l,…,4}; where *y* is the number of progeny/eggs produced by females after each cross, *f* is the treatment of origin of the females (monogamous or promiscuous, fixed effect), *m* is the treatment of origin of the males to which females were mated (monogamous or promiscuous, fixed effect) and *I* is their interaction. The interaction term was subsequently dropped, because it was not significant in any trial and did not improve the fit of the models (P > 0.25 for all models).

Microarray data were analysed using the BioConductor suite of packages (Gentleman *et al.*, 2004) in R. To pre-process the raw expression data, we used the standard RMA (Robust Multichip Average) algorithm (Irizarry *et al.*, 2003) implemented in the Affy package (Gautier *et al.*, 2004). After pre-processing the resulting dataset was filtered to exclude features according the following criteria: (i) probe sets without an Entrez Gene ID annotation, (ii) Affymetrix quality control probe sets, (iii) if multiple probe sets mapped to the same Entrez Gene ID, only the probe set with the highest coefficient of variation was retained. Out of the original 18952 features, the filtering step removed 6380 probe sets, while 12572 probe sets, corresponding to as many known genes, were retained for the statistical analyses.

Significance of differential expression was assessed using the package Limma (Linear Models for Microarray Data; Smyth, 2005). A model matrix was designed to fit a parameter for every combination of replicate population of origin of females (n = 8) and population of origin of males to which females were mated (n = 8), for a total of 16 parameters. An additional random effect with two levels was fitted to control for the batch effect, and estimated borrowing information between features, by constraining the within-block correlations to be equal across features and by using empirical Bayes methods to moderate the standard deviations (Smyth, 2005). A contrast matrix was designed to obtain the contrasts of interest: the main effect of treatment of origin of females, the main effect of treatment of origin of males to which females were mated, and their interaction. All the resulting P values were corrected for multiple testing to obtain a maximum false discovery rate of 5% (FDR; Benjamini & Hochberg, 1995; corrected P < 0.05).

We used a mean-rank gene set enrichment test (MR-GSE test, implemented in LIMMA; Michaud *et al.*, 2008) to test whether the sets of up-regulated or down-regulated significant transcripts showed a tendency to be up-or down-regulated after mating, using the t-values from a contrast between virgin and mated females from the same population, from a previously published study (Innocenti & Morrow, 2009). We defined as “Virgin-like’ the subset of transcripts for which the expression in the monogamous female is lower than the promiscuous females if the gene is up-regulated by mating, or higher if the gene is down-regulated by mating; in other words, genes whose profile is more similar to a virgin fly. ‘Mated-like’ genes are the complimentary set of genes. A MR-GSE test was also used to test whether the set of significant transcripts showed a tendency to be associated with female fitness, using the t-values from a previously published study (Innocenti & Morrow, 2010) on the same population.

Among the genes found to be differentially expressed, we identified transcriptional modules of correlated expression across-tissues using the Hopach package (Hierarchical Ordererd Partitioning and Collapsing Hybrid; van der Laan & Pollard, 2003). We computed a distance matrix using the pairwise correlations *r_ij_* between the expression of the significant transcripts across different tissues of *D. melanogaster.* The tissue-specific expression data were produced by the FlyAtlas team (Chintapalli *et al.*, 2007), available on the Gene Expression Omnibus (GEO) database with accession number GSE7763, and normalized according to a method described elsewhere (Innocenti & Morrow, 2010). The clustering algorithm built a hierarchical tree by recursively partitioning or collapsing clusters at each level, using MSS (Median Split Silhouette) criteria to identify the level of the tree with maximally homogeneous clusters (van der Laan & Pollard, 2003).

We selected the modules containing more than 50 genes (the clusters more likely to provide biologically meaningful within-module summary statistics) and analyzed them to identify whether they showed: (i) association with genes involved in female post-mating response (see above); (ii) association with female fitness (see above); (iii) non random chromosomal distribution; (iv) over-represented Gene Ontology categories; (v) tissue enrichment or specificity. Non-random chromosomal distribution was assessed with a Fisher’s exact test on the expected and observed number of genes on each chromosome (P < 0.01). To identify GO categories enriched for particular subsets of transcripts, we used a hypergeometric test for over-representation (P < 0.01, GOSTATS package; Falcon & Gentleman, 2007). We identified tissue-enriched or tissue-specific transcripts using data from the FlyAtlas database (described above). The tissue specific expression levels for the list of transcripts in each module were obtained, and the modules were tested for over-abundance of genes of interest in a target tissue using a one-tailed Fisher’s exact test. All the reported P values were Bonferroni corrected for testing on multiple tissues (P < 0.01, n = 17).

## Results

### Body size

After 30 generations of experimental evolution the body sizes of individuals from the two selection regimes remained virtually unchanged, with males and females of promiscuous lines having approximately 1% smaller wings than those in monogamous lines (mating system effect: *F*_3,12_ = 1.84, P = 0.200; Fig. 1).

**Figure 1.**
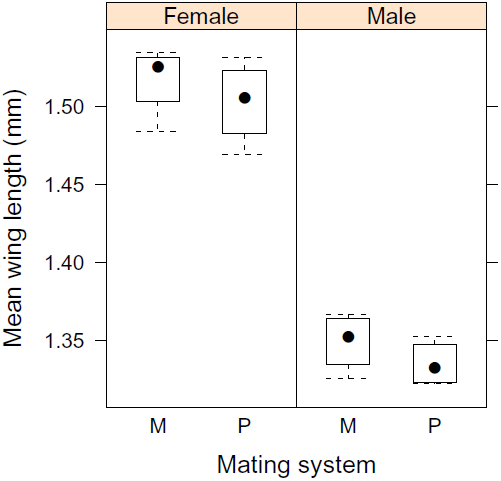
*Body size.* Wing length of females and males evolved under monogamous (M) and promiscuous (P) mating systems.

### Female fecundity

The reproductive output of promiscuous females was greater than those of monogamous females, regardless of the males they were mated with (Fig. 2; see Material and Methods). This difference was significant when measured as number of eggs laid in a 18 h period, corresponding to the oviposition period in every generation of experimental evolution (as well as the ancestral population), both after a single mating with a male (*F*_1,13_ = 4.89, P = 0.046), or being continuously exposed to males (*F*_1,13_ = 11.32, P = 0.005). When measured as number of adult progeny emerging from eggs laid during a period of 4 days, this difference was significant only after a single mating (*F*_1,13_ = 6.06, P = 0.029), but not when females where continuously exposed to males (*F*_1,13_ = 2.93, P = 0.110), although the effect sizes were comparable for direction and magnitude (Table 1).

**Figure 2.**
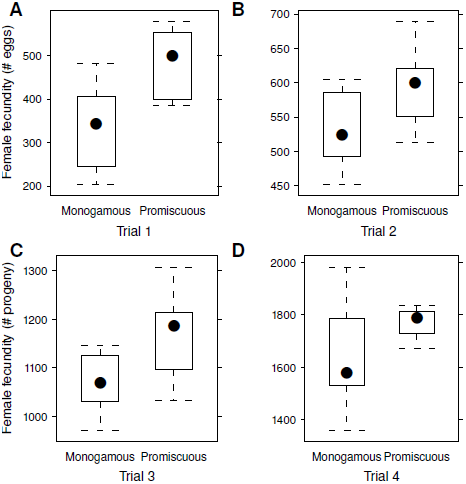
*Female fecundity.* Reproductive output of females evolved under monogamous and promiscuous selection regimes after mating once (panels B and C) or being continuously exposed to males (panels A and D), during the normal reproductive window (panel A and B) or during a longer interval (4 days; panel C and D).

**Table 1.**
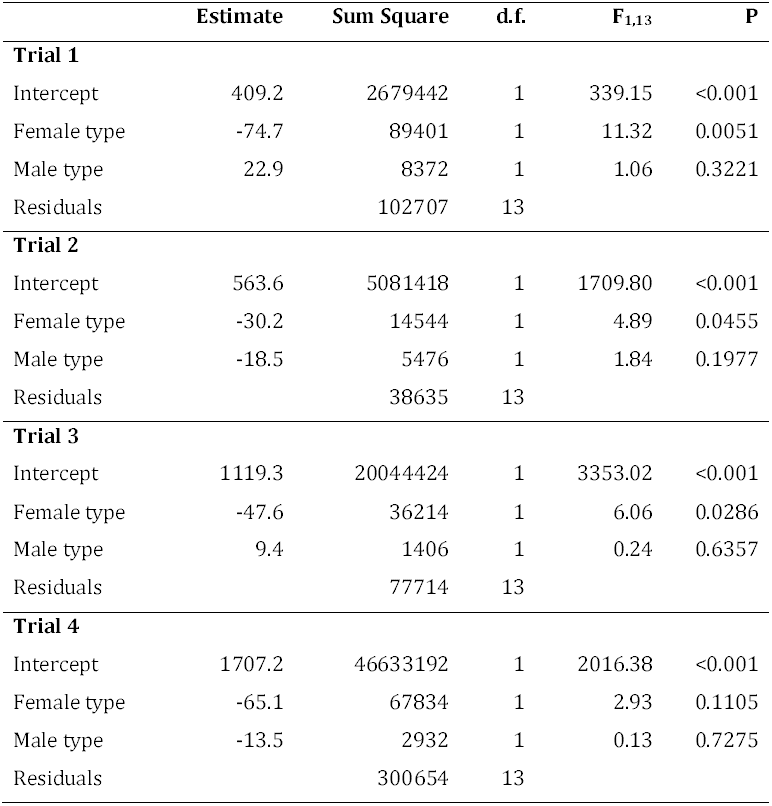
Linear model results on female fecundity. *Trial 1*: Females continuously exposed to males, 18 h oviposition period; *Trial 2*: Females mated once, 18 h oviposition period; *Trial 3*: Females mated once, 4 days oviposition period; *Trial 4*: Females continuously exposed to males, 4 days oviposition period.

|  | **Estimate** | **Sum Square** | **d.f.** | **F<sub>1,13</sub>** | **P** |
| --- | --- | --- | --- | --- | --- |
| **Trial 1** |  |  |  |  |  |
| Intercept | 409.2 | 2679442 | 1 | 339.15 | <0.001 |
| Female type | -74.7 | 89401 | 1 | 11.32 | 0.0051 |
| Male type | 22.9 | 8372 | 1 | 1.06 | 0.3221 |
| Residuals |  | 102707 | 13 |  |  |
| **Trial 2** |  |  |  |  |  |
| Intercept | 563.6 | 5081418 | 1 | 1709.80 | <0.001 |
| Female type | -30.2 | 14544 | 1 | 4.89 | 0.0455 |
| Male type | -18.5 | 5476 | 1 | 1.84 | 0.1977 |
| Residuals |  | 38635 | 13 |  |  |
| **Trial 3** |  |  |  |  |  |
| Intercept | 1119.3 | 20044424 | 1 | 3353.02 | <0.001 |
| Female type | -47.6 | 36214 | 1 | 6.06 | 0.0286 |
| Male type | 9.4 | 1406 | 1 | 0.24 | 0.6357 |
| Residuals |  | 77714 | 13 |  |  |
| **Trial 4** |  |  |  |  |  |
| Intercept | 1707.2 | 46633192 | 1 | 2016.38 | <0.001 |
| Female type | -65.1 | 67834 | 1 | 2.93 | 0.1105 |
| Male type | -13.5 | 2932 | 1 | 0.13 | 0.7275 |
| Residuals |  | 300654 | 13 |  |  |

### Reversed selection lines

As described in the Material and Methods, the reversed selection lines (MP) were established after the monogamous and promiscuous populations had already undergone 95 generations of selection. All three treatments were then run for a further 25 generations prior to the final assays of fecundity being performed. Despite this substantial additional period of experimental evolution, relative to the first round of assays (a further 90 generations), the patterns of reproductive output between monogamous and promiscuous females was remarkably similar at these two time points (M vs P differences: G30 45%; G120 36%), indicating that the majority of phenotypic evolution had occurred within the first 30 generations (posthoc Tukey HSD: M vs P, *t* = 2.716, P = 0.0279; Fig 3). The phenotypic change in reproductive output of females from the MP populations following 25 generations of reversed selection was smaller. It did however indicate a reversal in reproductive output had occurred; posthoc tests showed that the reproductive output of females from the MP treatment was intermediate to both monogamous and promiscuous treatments (Tukey HSD: M vs MP, *t* = 0.913, P = 0.6365; MP vs P, *t* = 1.804, P = 0.1850; Fig. 3).

**Figure 3.**
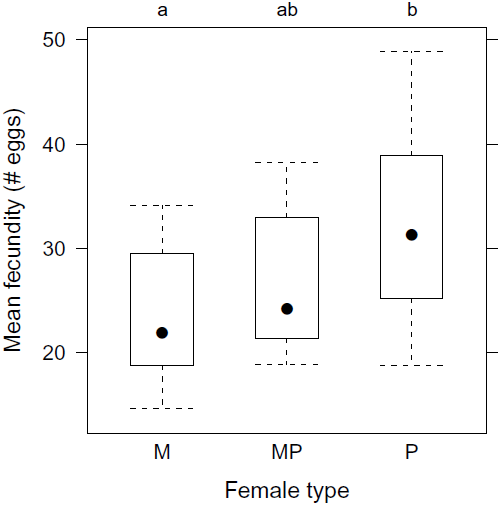
*Reversed selection.* Reproductive output of females evolved under monogamous (M), monogamous then promiscuous (MP) and promiscuous (P) selection regimes. Results of the posthoc analysis are given above the plotting frame, letters not shared indicate treatments that show statistical significant differences (see Results for details).

### Gene expression profiles

After 46 generations of selection, we tested the difference in female genome-wide post-mating response, by measuring gene expression in adult *D. melanogaster* females evolved under different sexual selection regimes (monogamous and promiscuous). After multiple testing correction, monogamous and promiscuous females showed a significant difference in the expression of 1141 transcripts (≈9% of the transcripts analyzed, at 5% F.D.R.), while male type and the interaction of female type and male type did not significantly affect the post-mating expression patterns. Among the differentially expressed transcripts, 438 were up-regulated and 703 down-regulated in monogamous females (Binomial test: ratio = 0.38, P < 0.0001).

We compared the expression profile of these transcripts with the female post-mating response characteristic to the ancestral population (Innocenti & Morrow, 2009), and found that the expression level of 728 genes is altered in monogamous females to a lesser extent after mating, compared to promiscuous females (hereafter ‘virgin-like’, see Material and Methods), while the post-mating reaction of the remaining 413 genes is altered to a higher extent in monogamous females compared to promiscuous females (hereafter ‘mated-like’), and their proportion is higher than expected by chance (Binomial test: ratio = 0.64, P < 0.0001). In general, genes that are down-regulated in monogamous vs. promiscuous mated females tend to be switched on (up-regulated) by mating (two-tailed MR-GSE test: P < 0.0001, Fig. 4A) and genes that are up-regulated in monogamous vs. promiscuous mated females tend to be switched off (down-regulated) by mating (two-tailed MR-GSE test: P < 0.0001, Fig. 4A). We also used previously published and independently derived data (Innocenti & Morrow, 2010) to test the relationship between the significant transcripts identified in this study and female fecundity, and found them to be over-represented among genes strongly associated with female fitness (irrespective of the sign of the association; MR-GSE test, P < 0.0001, Fig. 4B).

**Figure 4.**
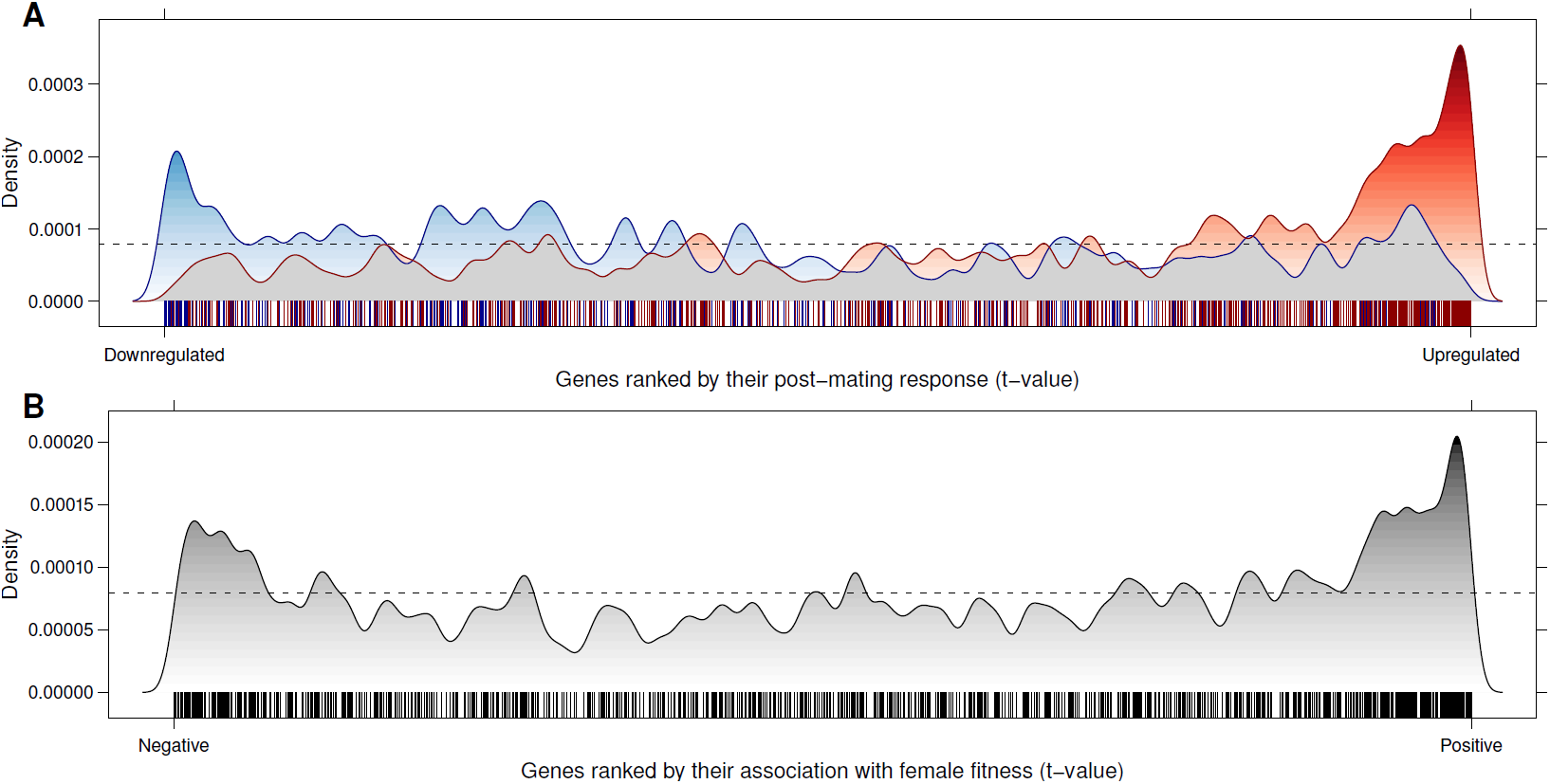
*Association with post-mating response and female fitness.* (A) Density distribution of significant up-regulated (blue) and down-regulated (red) transcripts along all the tested genes, ranked according to their post-mating reaction (data from a previously published study on the same population; Innocenti and Morrow, 2009). (B) Density distribution of the significant transcripts along all the tested genes, ranked by the t-value of their association with female fitness (data from a previously published study on the same population; Innocenti and Morrow, 2010).

In order to identify clusters of transcripts co-expressed in one or more tissues, and hence possibly involved in similar biological function, we calculated modules of correlated expression among the significant transcripts using data from the FlyAtlas database (Chintapalli *et al.*, 2007). Among them, we selected the 7 clusters containing more than 50 genes (Fig. 5), which represented about 75% of the significant transcripts, and evaluated their post-mating expression profile in comparison to the ancestral population, their tissue specificity, chromosomal distribution and over-presentation among Gene Ontology categories (see Supplementary Information).

**Figure 5.**
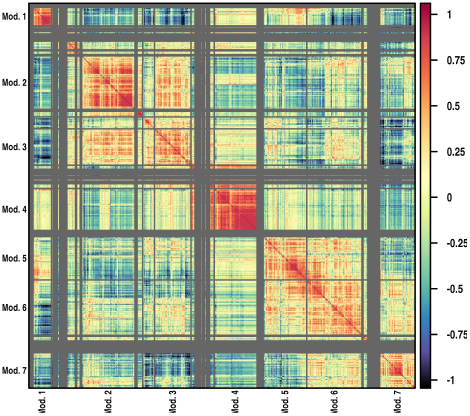
*Transcriptional modules.* Level-plot representing the matrix of pairwise correlation for the expression of the 1141 significant transcripts across tissues of *D. melanogaster* (data from Chintapalli *et al.*, 2007). The correlation matrix has been used to compute modules of correlated expression (separated by grey lines). The 7 modules containing more than 50 genes are labeled.

Module 1 contains genes highly specific for the male gonads, showing little or no expression in other tissues (Fig. S1D,E). Overall, they do not tend to be perturbed by mating (Fig. S1A). Over-represented Gene Ontology (GO) terms indicate that the activity of a portion of these genes is linked to mitochondrial cellular components (cellular respiration, electron transport, Table SI).

Module 2 is a large cluster of genes active in the majority of the tissue types (but generally not in the gonads, Fig. S2E), and significantly over-expressed in the head, eyes, carcass, fat body, heart and spermatheca. These transcripts are subject to changes in expression levels after mating (Table 2), with mated monogamous flies showing a more virgin-like expression profile for these genes (Fig. S2A). They are chiefly involved in enzymatic metabolic activity (oxidation reduction, proteolysis, Table S2). The left arm of chromosome 2 is enriched for this set of genes (Fisher exact test: 0dds-ratio = 1.49, P = 0.005).

**Table 2.** Summary description of the main modules. ‘*Size*’: Number of significant genes in each module (n > 50). ‘*Tissue*’: tissues in which the transcripts in each module are significantly over-expressed compared to the whole body (data from Chintapalli *et al.*, 2007). ‘*Up*’ (‘*Down*’) is the subset of up-regulated (down-regulated) transcripts in the module. In parentheses is indicated whether the subset tend to be up-regulated or down-regulated after mating, or a mix of the two (mixed); P value from a MR-GSE test; see also Figs. S1A-S7A, (data from Innocenti & Morrow, 2009). ‘*Fitness assoc.*’ indicates whether the genes in the module are over-represented among the genes found to be associated with female fitness (MR-GSE test, data from Innocenti & Morrow, 2010).

| Module | Size | Tissue | Up | Down | Fitness assoc. | GO terms |
| --- | --- | --- | --- | --- | --- | --- |
| 1 | 55 | Testes | 24 (n.s.) | 31 (n.s.) | Yes (0.016) | Table S1 |
| 2 | 158 | Carcass, Head, Eyes, Heart, Fat body,<br>Spermatheca | 52 (mixed, <0.001) | 106 (up, <0.001) | Yes (<0.001) | Table S2 |
| 3 | 108 | Crop, Mid gut, Hind gut, Heart, Fat body,<br>Spermatheca | 42 (mixed, 0.007) | 66 (n.s.) | Yes (<0.001) | Table S3 |
| 4 | 124 | Mid gut | 33 (mixed, 0.002) | 91 (up, <0.001) | Yes (<0.001) | Table S4 |
| 5 | 129 | Ovary | 83 (down, <0.001) | 46 (n.s.) | Yes (0.006) | Table S5 |
| 6 | 164 | Brain, Ovary | 113 (n.s.) | 51 (down, 0.002) | Yes (0.014) | Table S6 |
| 7 | 105 | Brain, Eyes, Head, Thoracic ganglion | 25 (n.s.) | 80 (up, <0.001) | Yes (0.047) | Table S7 |

The activity of genes clustered in module 3 is very similar to those of module 2: these transcripts are active ubiquitously in the fruit fly tissues (Fig. S3D,E) and the up-regulated subset is significantly enriched among the set of genes which respond to mating (Table 2). Although not significant under our cut-off, a higher than expected proportion of these genes lies on chromosome 2L (Odds-ratio = 1.47, P = 0.045). GO terms associated with these genes include, again, strong cytoplasmic enzymatic activity (oxidation reduction, catalytic activity, Table S3).

Module 4 presents the most distinctive and peculiar patterns. The majority of these genes are down-regulated in the monogamous treatment (91 out of 124, Table 2) and tend to be strongly affected by mating and distinctly more virgin-like in monogamous females (Fig. S4A). These transcripts are consistently highly expressed in the midgut, but relatively silent in all the other tissues (Fig. S4D,E), and most of their activity is linked to metabolic processes, mainly peptidase and hydrolase activity (Table S4). The distribution on the chromosomes is significantly skewed towards the right arm of chromosome 2 (Odds-ratio = 1.49, P = 0.009).

Modules 5 and 6 show highest relative expression levels in the ovaries, although the transcripts are also active at slightly lower levels in every other tissue. These genes tend to be overall weakly down-regulated after mating (Figs. S5A and S6A). Module 5 showed relative virgin-like expression in monogamous females compared to promiscuous females (Table 2). Perhaps unsurprisingly, sexual reproduction and female gamete generation were among the most enriched biological processes, while the same sets of genes were linked to nucleic acid and protein binding molecular functions (Table S5). Module 6, also significantly over-expressed in the brain, showed enrichment for biological processes such as behaviour and signaling processes (Table S6).

Module 7 contains genes significantly more active in neural tissues: brain, thoracic ganglion, head and eyes (Fig. S7D,E). They are mostly down-regulated (Table 2) in monogamous females, but tend to be up-regulated after mating (virgin-like in monogamous females, Fig. S7A).

All modules tend to be associated with female fitness, with transcripts in modules 2, 3 and 4 showing the strongest association (Table 2).

## Discussion

In our study, we experimentally manipulated the mating system in replicate populations of *D. melanogaster*, by removing sexual selection, with the aim of testing differences in short term post-mating reaction of females evolved under different mating strategies. We showed that monogamous females suffer decreased fecundity, regardless of the type of male they were mated with, or whether mated once or continuously exposed to males. We also showed that monogamous females could recover some of this loss in fecundity if the selection pressure was reversed experimentally. Previously, Holland and Rice (1999) removed sexual selection in experimental lines from the same population (LH_M_) by manipulating sex ratio, and found that (i) monogamous females showed higher ‘net reproductive rate’ (female fecundity and offspring survival) than controls when mated with males from their own populations, and (ii) monogamous females showed lower fecundity than controls when mated once to ancestral (promiscuous) males (Holland & Rice, 1999). Monogamous males, in turn, evolved decreased courtship rate. Our experiment employed a different design, which allowed mass mating (mate choice and pre-copulatory intra-sexual competition) but a single mating event in the monogamous treatment, in order to leave selection on courtship rate unaffected. Our results showed no effect of male type on female fecundity (and consequently no interaction between male and female type), which rules out the possibility that males evolved decreased courtship intensity or a less harmful ejaculate. It is thus unlikely that the decrease in fecundity of monogamous females reflects a selective pressure towards less ‘resistant’ females. On the other hand, the experimental treatment removed continuous male harassment in the monogamous environment and decreased population density during selection. Relaxed selection on resistance to male harassment may have allowed the accumulation of deleterious mutations or recombination of extant genetic variation with sub-optimal epistatic effects which resulted in overall decrease in mean female fitness in the monogamous environment. The decline in female fitness in monogamous lines is not likely to be due to simple differences in population size and subsequent inbreeding, since not only does theoretical and previous empirical work indicate that n = 96 is above the threshold for drift decay (see Morrow *et al.*, 2008), our reversed experimental evolution treatment showed that the significant differences in fecundity between monogamous and promiscuous females disappeared when monogamous females experienced a reintroduction of a promiscuous mating system. Such a response would not occur if monogamous populations had simply become bottlenecked. This is further supported by the minimal differences in body size seen between monogamous and promiscuous treatments, a trait that could be sensitive to inbreeding.

The results of our genome-wide expression analysis confirmed a significant difference between post-mating reaction between monogamous and promiscuous females, while the evolutionary history of the males to which they were mated did not influence their expression profiles. The genes that evolved to respond differently to mating accounted for around 9% of the transcriptome tested. When comparing transcriptional changes which occur when a female switches between the virgin and mated status with differential expression between mated monogamous and promiscuous females, it is clear that genes which are up-regulated by mating tend to be down-regulated in monogamous vs. promiscuous females (and those down-regulated by mating tend to be up-regulated in monogamous vs promiscuous females; Fig. 4A), i.e. monogamous females generally show a more ‘virgin-like’ expression profile. Similarly, significant genes tend to be over-represented among candidate genes known to be associated with female fitness. Taken together, these two lines of evidence can be interpreted as strong support for the phenotypic results showing decreased female fecundity: monogamous females seem to exhibit a weaker post-mating reaction, both in terms of the extent to which genes are expressed and how many eggs they lay.

When analysing and partitioning these genes according to the tissue where they are predominantly active, we can identify 4 broad categories: transcripts active in i) the midgut, ii) the ovaries, iii) neural tissues and iv) a wide range of tissues. The midgut (module 3, Table 2) provides the strongest and clearest signal, (Fig. S4E), and genes active in this tissue are mainly linked to enzymatic activity (Table S4). Such genes are usually activated by mating and show a decreased response in monogamous females (Fig. S4A). Significant genes in the ovaries (module 5 and 6) are involved in gamete production, while in the neural tissues they regulate signaling processes and transmembrane transport activity (Table S5, S6). The last category (module 2 and 3) contains genes expressed in a diverse array of tissues (Table 2), although known to show overall very high correlation for expression (Innocenti & Morrow, 2010). The Gene Ontology categories involved (oxidoreductase activity, lipid and sugars storage/metabolism, Table S2, S3) seem to indicate a predominant function in energy production and resource consumption. An additional, small category points to transcripts mostly active in the testes, and its interpretation is problematic, given the sex-limited nature of this tissue. This set of genes, however, which are not involved in a normal post-mating reaction (Fig. S1A), could be selected due to pleiotropic activity in other tissues, or exhibit non-random segregation (e.g. linkage disequilibrium) with selected transcripts.

In *D. melanogaster*, transfer of the ejaculate, and in particular some seminal fluid components (e.g. Sex Peptide), radically transforms female behaviour and physiology, leading to increased egg production (Gillott, 2003), decreased receptivity (Chapman *et al.*, 2003) and increased feeding behaviour (Carvalho *et al.*, 2006), and has profound effects on gene expression, especially gene products affecting metabolic rate (Innocenti & Morrow, 2009). Moreover, at least part of these physiological changes are mediated by the nervous system (Hasemeyer *et al.*, 2009). Together this diverse range of evidence confirms that monogamous females are less fecund than promiscuous females because they exhibit a weaker post-mating response. The origin of this difference in response might reflect a lower ‘genetic quality’ of monogamous females arising from a relaxed selection on female resistance to male harassment or from relaxed post-copulatory selection, which can directly affect female ability to induce a physiological response to mating, or be mediated by an overall weaker condition. In this respect, Fricke *et al.* (2010) recently showed how nutritional status of females determines their response to the sex peptide and influences fecundity, with high food diets being associated with increased egg production, raising the hypothesis that female response to mating can be environmental or condition dependent.

This study, in combination with the independent characterization of post-mating expression profiles in females and the relationship between transcript abundance and female fitness in the ancestral population, provides a robust list of candidate genes associated with changes in female fecundity caused by evolution under different sexual selective pressures. Given the close agreement between what is already known about the effects of the male ejaculate on females and their fitness in *D. melanogaster*, with the general patterns of tissue specificity and biological processes identified here, these data provide a clear indication as to which genes are the targets of post-mating sexual selection in this promiscuously mating population.

## Acknowledgments

We thank Jessica Abbott, Tim Connallon and Björn Rogell for comments on the manuscript and the Uppsala Array Platform. Funding was provided by the Swedish Research Council (Grant #2008-5533 and #2011-3701), the European Research Council (Grant #280632) and a Royal Society University Research Fellowship to EHM.

